# Control of asparagine homeostasis in *Bacillus subtilis*: Identification of promiscuous amino acid importers and exporters

**DOI:** 10.1101/2023.12.05.570048

**Authors:** Janek Meißner, Manuel Königshof, Katrin Wrede, Robert Warneke, Mohammad Saba Yousef Mardoukhi, Fabian M. Commichau, Jörg Stülke

## Abstract

Amino acids are the main building block for proteins. The Gram-positive model bacterium *B. subtilis* is able to import all proteinogenic amino acids from the environment as well as to synthesize them. However, the players involved in the acquisition of asparagine have not yet been identified for this bacterium. In this work, we used D-asparagine as a toxic analog of L-asparagine to identify asparagine transporters. This revealed that D-but not L-asparagine is taken up by the malate/lactate antiporter MleN. Specific strains that are sensitive to the presence of L-asparagine due to the lack of the second messenger cyclic di-AMP or due to the intracellular accumulation of this amino acid were used to isolate and characterize suppressor mutants that were resistant to the presence of otherwise growth-inhibiting concentrations of L-asparagine. These screens identified the broad-spectrum amino acid importers AimA and BcaP as responsible for the acquisition of L-asparagine. The amino acid exporter AzlCD allows detoxification of L-asparagine in addition to 4-azaleucine and histidine. This work supports the idea that amino acids are often transported by promiscuous importers and exporters. However, our work also shows that even stereo-enantiomeric amino acids do not necessarily use the same transport systems.

**IMPORTANCE:** Transport of amino acid is a poorly studied function in many bacteria, including the model organism *Bacillus subtilis*. The identification of transporters is hampered by the redundancy of transport systems for most amino acids as well as by the poor specificity of the transporters. Here, we apply several strategies to use the growth-inhibitive effect of many amino acids under defined conditions to isolate suppressor mutants that exhibit either reduced uptake or enhanced export of asparagine, resulting in the identification of uptake and export systems for L-asparagine. The approaches used here may be useful for the identification of transporters for other amino acids both in *B. subtilis* and other bacteria as well.

## INTRODUCTION

Every living cell is dependent on a steadily available pool of amino acids to maintain constant protein biosynthesis. The Gram-positive model organism *Bacillus subtilis* is able to synthesize all twenty proteinogenic amino acids, while using glucose and ammonium as the only sources of carbon and nitrogen, respectively. Amino acid homeostasis can be managed by the interplay of uptake, biosynthesis, degradation, export, and consumption for multiple metabolic purposes. The pathways of amino acid biosynthesis and degradation are well understood for *B. subtilis* and many other bacteria (1). In contrast, the transport of amino acids has recently been identified as a topic that requires much more attention as there are still several amino acids, among them asparagine, glycine, phenylalanine, and tyrosine for which no transporter has been identified so far. On the other hand, no substrate has so far been identified for many potential amino acid transporters (2). These gaps in our knowledge need to be filled to gain a comprehensive picture of amino acid homeostasis in *B. subtilis*, one of the most intensively studied organisms.

The identification of amino acid transporters is hampered by two problems: First, many amino acids are taken up by multiple transporters, and the deletion of one or even several candidate genes may not give rise to a clear phenotype. On the other hand, many amino acid transporters are promiscuous, *i.e.* they can transport multiple amino acids. Both issues have also been observed in *B. subtilis*. For example, glutamate can be transported by the amino acid permeases GltT and AimA (3, 4). For the branched-chain amino acids isoleucine and valine, BcaP, BrnQ, and BraB have been identified as permeases (5). Similarly, three permeases, *i.e.* PutP, OpuE, and GabP, are involved in the acquisition of proline (6, 7, 8). Concerning transporter promiscuity, the amino acid transporter AimA is involved in the acquisition of serine and glutamate, whereas BcaP, which was originally identified as a transporter for branched-chain amino acids, also transports threonine and serine (3, 4, 5, 9). Similarly, the bipartite amino acid exporter AzlCD can export a branched-chain amino acid analog, 4-azaleucine as well as histidine (10, 11). Amino acid transporters can be identified by expressing them in established mutants that lack transporters for particular amino acids and that are therefore unable to utilize the target amino acid as a carbon and/or nitrogen source. In this way, by complementation of *Escherichia coli* mutants, AimA could be shown to transport both serine and glutamate (3, 4). Alternatively, strains that are auxotrophic for a given amino acid can be tested for the effect of the inactivation of putative transporter gene(s). This strategy was used to identify the alanine permease AlaP (12). Finally, the identification of transport systems can profit from the fact that some amino acids inhibit the growth of bacteria. For threonine and serine, this growth inhibition has been described in both *E. coli* and *B. subtilis* (13, 14, 15, 16). Some amino acids become toxic if they accumulate due to the lack of the corresponding catabolic enzymes. Similarly, strains lacking the essential second messenger c-di-AMP are highly sensitive to multiple amino acids (4, 11, 17), probably resulting from the formation of glutamate which drastically increases the affinity of the the potassium channel KtrCD for potassium thus supporting potassium uptake which is toxic for such mutants (18, 19, 20). Finally, many D-amino acids are toxic for the cells, and toxic analogs exist for many amino acids, such as 4-azaleucine (21, 22). Irrespective of the context, amino acid toxicity has been proven a powerful tool to identify amino acid transporters, particularly by isolating suppressor mutants that are viable in the presence of otherwise toxic amino acids, and the identification of the responsible mutations. In this way, transporters for glutamate, proline, serine, and threonine have been discovered (3, 4, 5, 8, 9).

We have a long-standing interest in amino acid metabolism and transport in *B. subtilis* (23, 24, 25, 26, 27, 28, 29, 30). In this work, we have addressed the transport of asparagine. Asparagine is one of the amino acids for which no importer has been identified so far in *B. subtillis*. Asparagine is synthesized from aspartate, via the asparagine synthetases AsnO, AsnH and AsnB. These enzymes all use glutamine as the amide group donor to form asparagine from aspartate. The degradation of asparagine via the asparaginases AnsA and AnsZ leads back to aspartate. Aspartate then feeds into the citric acid cycle via the aspartase AnsB. The asparaginase and aspartase are encoded in the asparagine-induced *ansAB* operon (31), which also includes *mleN*, coding for a malate/lactate antiporter (32), as well as the *mleA* gene, which encodes the malic enzyme MleA that decarboxylates malate to pyruvate (33, 34, 35).

Using a variety of suppressor screens, we provide evidence that the malate/lactate antiporter MleN is responsible for the uptake of D-asparagine whereas AimA and BcaP are responsible for the uptake of L-asparagine. The latter amino acid can also be exported by mutational activation of the AzlCD amino acid exporter.

## RESULTS

### *D*-Asn is toxic for *B. subtilis*, and mutations in the *mleN* gene overcome this toxicity

Several D-amino acids are harmful to *B. subtilis*, as they exert toxic effects during protein synthesis (21). We tested the effect of D-Leu, D-Arg, D-Gln, D-Asp and D-Asn on the growth of *B. subtilis*. Of these D-amino acids only D-Asn and D-Gln showed toxic effects (the results with D-Asn are shown in Fig. 1). The wild type strain *B. subtilis* 168 was unable to grow at D-Asn concentrations above 5 mM. We isolated stable suppressor mutants that were able to grow in the presence of D-Asn. We isolated four suppressor mutants and subjected two of them to whole genome sequencing. The data revealed an identical single mutation in the malate/lactate antiporter MleN (32) in both mutants, causing the insertion of two base pairs, resulting in a premature stop codon, and thus likely in the inactivation of the transporter (see Table 1). We then sequenced the *mleN* region of the two remaining suppressor mutants, and found that the *mleN* gene was mutated there as well. Interestingly, all mutants had insertions of one or two base pairs in the *mleN* coding region that caused frame shifts in the N-terminal half of the protein.

**FIG 1.**
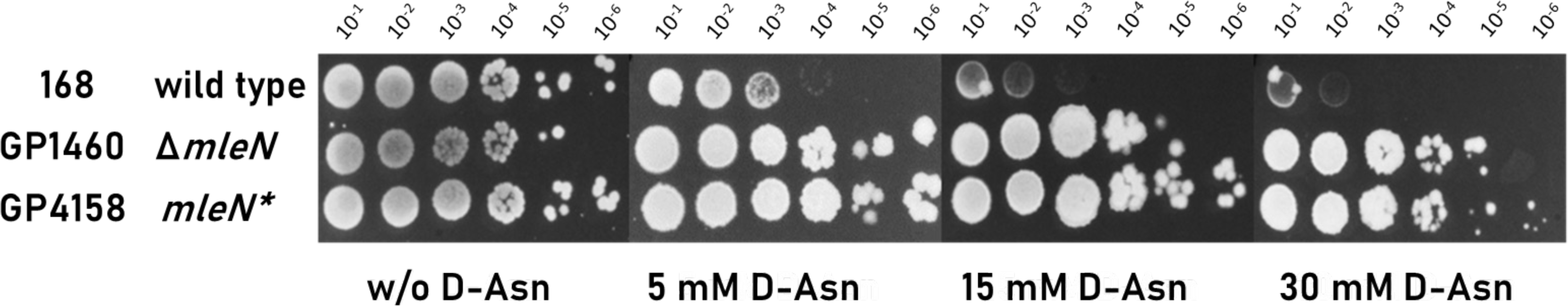
The *mleN* mutation confers resistance to D-asparagine stress. Sensitivity of wild type *B. subtilis* (168) and the isogenic Δ*mleN* mutant (GP1460) as well as a suppressor mutant (GP4158) isolated in the presence of D-asparagine, which carries a mutation in *mleN*. The cells were grown in SM minimal medium to an OD_600_ of 1.0 and were then diluted 10-fold to create dilutions ranging from 10^-1^ to 10^-6^. The dilution series was dropped onto SM plates with and without D-asparagine (0, 5, 15 and 30 mM respectively). The plates were incubated at 37°C for 48 h.

**TABLE 1.** Identification of suppressor mutations^a^.

| Strain | Affected gene(s) | Mutation(s) |
| --- | --- | --- |
| *B. subtilis* 168 (wild type), selection on SM plates containing D-Asn (5 mM) |  |  |
| GP4158 | *mleN* | +GC at position 517, frameshift, premature stop codon |
| GP4159 | *mleN* | +GC at position 517, frameshift, premature stop codon |
| S3 | *mleN* | +T at position 691, frameshift, premature stop codon |
| S4 | *mleN* | +TG at pos 517, frameshift, premature stop codon |
| *B. subtilis* GP2222 ($\Delta dac$), selection on MSSM plates containing L-Asn (15 mM) | | |
| GP4269 | *ktrD* | –T at position 72, frameshift, premature stop codon |
| GP4270 | *ktrD* | –CAGTCGG at position 935, frameshift, premature stop codon |
| S3 | *ktrD* | *ktrD*<sub>G314W</sub> |
| S4 | *ktrD* | *ktrD*<sub>G314W</sub> |
| *B. subtilis* GP4197 ($\Delta ansAB \Delta ansZ$), selection on C glucose plates containing L-Asn | | |
| GP4230 (5mM L-Asn) | *azlB* | *azlB*<sub>N24S</sub> |
| GP4231 (10mM L-Asn) | *azlB* | +CATTAAATG at position 37, frameshift, premature stop codon |
| GP4232 (15mM L-Asn) | *azlB* | C<sub>388</sub> → T, Q130 → Stop codon, premature stop codon at pos. 130 |
| GP4233 (30mM L-Asn) | *azlB* | *azlB*<sub>D19G</sub> |
| *B. subtilis* GP4245 ($\Delta ansAB \Delta ansZ \Delta azlBCD \Delta aimA$), selection on C glucose plates containing L-Asn (isolated on disc diffusion plates, see Fig. 4) | | |
| S1 | *bcaP* | C<sub>982</sub> → T, Q328 → Stop codon, premature stop codon at pos. |
|  |  | 328 |
| S2 | *bcaP* | C<sub>982</sub> → T, Q328 → Stop codon, premature stop codon at pos.<br>328 |
| GP4251 | *bcaP*<br>*ydeC* | *bcaP*<sub>A68E</sub><br>*ydeC*<sub>H43Q</sub> |
| S4 | *bcaP* | C<sub>982</sub> → T, Q328 → Stop codon, premature stop codon at pos.<br>328 |
| S5 | *bcaP* | –50 bp at position 153, frameshift, premature stop codon |
| S6 | *bcaP* | –50 bp at position 153, frameshift, premature stop codon |
| GP4252 | *bcaP* | C<sub>982</sub> → T, Q328 → Stop codon, premature stop codon at pos.<br>328 |
| S8 | *bcaP* | BcaP<sub>V92M</sub> |
| *B. subtilis* BP269 (SP1 ΔansAB), selection on LB plates containing L-Asn (19 mM) | | |
| BP391 | *aimA*<br>*yetL* | Duplication of 191CTA193, insertion of Thr after Ser-64<br>YetL<sub>A96E</sub> |
| BP392 | *aimA* | +G at position 213, frameshift, premature stop codon |

<sup>a</sup> For the complete genotypes of all strains, see Table 3. All gene designations are used in *SubtiWiki* (36), and the genes can be retrieved there.

We then examined the growth of an *mleN* deletion strain (GP1460), compared to the wild type strain (168) and one of the suppressors (GP4158), in the presence of increasing amounts of D-Asn (see Fig. 1). While low amounts of D-Asn (up to 5 mM) could be tolerated by the cells, higher amounts led to cell death and suppressor formation in the wild type strain. Both the suppressor mutant GP4158 and the Δ*mleN* deletion mutant GP1460 were able to grow at up to 30 mM D-Asn. This result indicates that the truncation of the MleN protein in the suppressor mutant indeed results in an inactivation of the protein, and shows that this mutation is responsible for the suppression. The fact that the Δ*mleN* deletion led to full D-Asn resistance suggests that MleN can transport D-Asn in *B. subtilis*.

### Expression of the *mleN* gene

The putative D-Asn transporter MleN and the NAD^+^-dependent malate dehydrogenase MleA seem to be expressed in an L-asparagine-inducible operon with the asparaginase AnsA and the aspartase AnsB (31, 35). According to a large-scale transcriptome analysis with *B. subtilis* and the *Subti*Wiki database, there might be one promoter in front of *ansA* and a second promoter in front of *mleN* (35, 36). We wanted to verify this observation, as well as examine a possible induction of the operon by amino acids, particularly by the D- and L-enantiomers of asparagine and aspartate. For this purpose, we made use of strains that carry fusions of the regions upstream of the *ansA* and *mleN* genes to a promoterless *lacZ* reporter gene. The strains were grown in C glucose minimal medium in the absence or presence of casein hydrolysate, and the asparagine and aspartate enantiomers (see Table 2). In agreement with previous reports (31, 37), the *ansA* promoter was highly active in the presence of L-Asn, whereas only weak expression was observed under all other tested conditions. In contrast, only weak activities were observed for the *mleN* promoter. The observed readthrough in the *ansAB-mleNA* region (35) as well as the absence of a promoter in front of *mleN* suggest, that the four genes are co-expressed in the *ansAB-mleNA* operon, via the asparagine-inducible promoter in front of *ansA*. Due to the toxicity of D-Asn, we were unable to assay promoter activities in the wild type background. To circumvent this problem, we transferred the *lacZ* fusion constructs into the strain lacking the D-asparagine importer MleN. In this case (GP4191), we observed a very weak increase of *ansA* promoter actvity which is, however, neglegible as compared to the observed induction of the promoter in the presence of L-Asn (Table 2) and in comparison to promoters of basic carbon metabolism (38). However, this activity seems to be sufficient to express the *mleN* gene to a level that allows uptake of D-Asn and concomitant intoxication of the *B. subtilis* cells.

**TABLE 2.** Activity of the putative *mleN* and *ansA* promoters.

| | | Units of β-galactosidase per μg of protein | | | | | |
| --- | --- | --- | --- | --- | --- | --- | --- |
|  |  | Addition to C-Glc minimal medium |  |  |  |  |  |
| Strain | Relevant genotype | None | CAA | D-Asn<sup>a</sup> | L-Asn<sup>a</sup> | D-Asp<sup>a</sup> | L-Asp<sup>a</sup> |
| GP829 | *mleN-lacZ* | 6 ± 1 | 11 ± 7 | NG<sup>b</sup> | 2 ± 0 | 56 ± 15 | 2 ± 0 |
| GP1155 | *ansA-lacZ* | 13 ± 1 | 52 ± 3 | NG | 1325 ± 247 | 18 ± 2 | 11 ± 2 |
| GP4191 | *ansA-lacZ* Δ*mleN* | 11 ± 2 | 47 ± 5 | 48 ± 23 | 1798 ± 83 | 30 ± 4 | 24 ± 7 |

<sup>a</sup> The amino acids were added to a concentration of 5 mM; <sup>b</sup> NG, no growth.

### The toxicity of L-Asn for a *Δ*dac mutant allows the identification of AimA as an L-Asn transporter

In some cases, the D- and L-enantiomers of an amino acids are taken up by the same transporter, as it is the case for D- and L-alanine that are both taken up by AlaP (12). This raised the question whether MleN might also be involved in the transport of L-Asn. This hypothesis is highly attractive since *mleN* is part of an operon involved in L-Asn utilization that is induced by this amino acid. The *B. subtilis* wild type strain 168 tolerates L-Asn in minimal medium (see Fig. 2). We made therefore use of the observation that a strain that is unable to produce c-di-AMP due to the deletion of the three genes encoding the diadenylate cyclases (Δ*dac*) is sensitive to glutamate and several other amino acids (4, 17). As shown in Fig. 2, the Δ*dac* mutant GP2222 is highly sensitive to growth inhibition by both D-Asn and L-Asn. To test the role of MleN in the uptake of L-Asn, we constructed the Δ*dac* Δ*mleN* mutant GP4177. Previous studies have shown that AimA is a major low affinity glutamate and serine importer in *B. subtilis* (3, 4). Thus, we also used the Δ*dac* Δ*aimA* mutant strain GP3054 (Fig. 2). The wild type as well as the single and double mutants were tested for growth in the absence and presence of D- and L-Asn. All strains grew well in the absence of any added amino acid. As expected, growth of the wild type strain 168 and the Δ*dac* mutant was inhibited In the presence of D-Asn whereas the deletion of *mleN* restored growth in both genetic backgrounds. In addition, the Δ*dac* Δ*aimA* double mutant was unable to grow in the presence of D-Asn, confirming that AimA does not contribute to the uptake of D-Asn. L-Asn was well tolerated by both the wild type and the Δ*mleN* mutant. In contrast, the Δ*dac* mutant GP2222 was unable to grow if L-Asn was present. The analysis of the double mutants revealed that only the deletion of *aimA* but not of *mleN* confers resistance to L-Asn. The drop dilution assay (see Fig. 2) shows that the deletion of *aimA* allows growth in the presence of L-Asn; however, growth was not restored to the full level as compared to the wild type strain, indicating that *B. subtilis* encodes additional L-Asn transporter(s). Taken together, our results indicate that MleN does not contribute to the uptake of L-asparagine, whereas AimA is able to transport this amino acid. This experiment demonstrates that the L- and D-forms of Asn, though structurally very similar, are not taken up by the same transporter.

**FIG 2.**
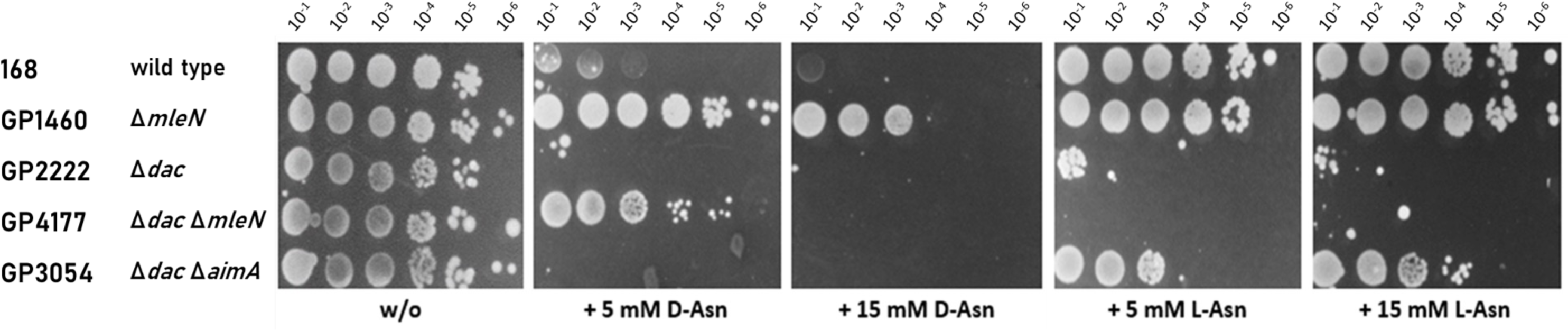
MleN is specific to D-asparagine import, while AimA is an L-asparagine importer. Sensitivity of wild type *B. subtilis* (168), the *mleN* mutant GP1460 and Δ*dac* mutants to D-Asn and L-Asn is compared on MSSM minimal medium. The cells were grown in MSSM minimal medium to an OD_600_ of 1.0 and were then diluted 10-fold to create dilutions ranging from 10^-1^ to 10^-6^. The dilution series was dropped onto MSSM plates with and without L-asparagine. The plates were incubated at 42°C for 48 h.

To get more insights into the sensitivity of the Δ*dac* mutant GP2222 to L-asparagine, we isolated suppressor mutants that were able to grow in the presence of 15 mM L-asparagine. Of four isolated mutants, two were subjected to whole-genome sequencing. Both mutants had frameshift mutations in the *ktrD* gene that result in the formation of truncated proteins (see Table 1). The *ktrD* alleles of the two remaining mutants were sequenced, and both had nucleotide substitutions that resulted in the replacement of Gly-314 by a Trp residue. The KtrD protein is the membrane subunit of the low affinity KtrCD potassium channel (39). It has been shown that the presence of glutamate results in high-affinity potassium uptake by KtrCD, resulting in intoxication of the Δ*dac* mutant (19). It is therefore possible that either L-asparagine itself can also activate KtrCD, or that KtrCD is activated by the aspartate formed upon asparagine degradation. The possible link between KtrCD and asparagine transport will be addressed elsewhere. The fact that we did not find suppressor mutations with inactivated amino acid transporter genes supports the idea that multiple transporters for L-Asn are present in *B. subtilis*.

### Identification of the AzlCD system as an potential exporter for L-Asn

Amino acids can be growth-inhibiting by themselves or due to the generation of harmful intermediates (16). To distinguish between these possibilities, we investigated the toxicity of L-asparagine in a strain that lacks the two asparaginases AnsA and AnsZ. This strain (GP4197) is sensitive to L-asparagine on minimal medium suggesting that the accumulation of L-asparagine itself is harmful for the cells (see Fig. 3). We observed suppressor mutant formation of the *ΔansAB ΔansZ* strain GP4197 after incubation at 42°C for 48h with 5, 10, 15 and 30 mM L-asparagine. We isolated four suppressor mutants, one for each of the tested concentrations of L-asparagine.

**FIG 3.**
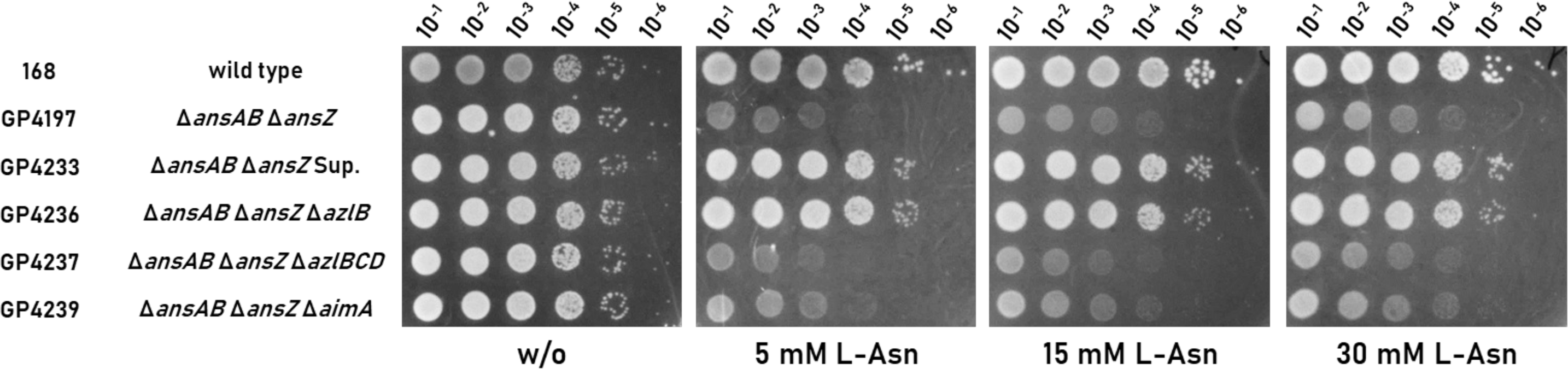
AzlCD provides resistance to L-asparagine when AzlB is mutated. Sensitivity of the Δ*ansAB* Δ*ansZ* (GP4197) double mutant is shown compared to the suppressor GP4233, isolated from L-Asn stress, as well as the knockout strains for the exporter AzlCD and the importer AimA. The cells were grown in C-Glc minimal medium to an OD_600_ of 1.0 and were then diluted 10-fold to create dilutions ranging from 10^-1^ to 10^-6^. The dilution series was dropped onto C-Glc plates with and without L-asparagine (0, 5, 15 and 30 mM respectively). The plates were incubated at 42°C for 48 h.

To identify the mutations responsible for the resistance to L-Asn, we performed whole genome sequencing for the four suppressor mutants. The sequencing results are summarized in Table 1. In each mutant, we found single mutations that all affected the transcriptional repressor AzlB, which controls the expression of the *azl* operon. Two of the mutations (in GP4231 and GP4232) result in the formation of truncated proteins (due to a frame shift or a point mutation that converts a codon for glutamine to a stop codon), and two mutations (in GP4230 and GP4233) result in AzlB proteins with amino acid substitutions. Two of the mutations, the eight base pair insertion of CATTAATG after nucleotide 37 (in GP4231) and the Asn24-Ser substitution (in GP4230) were already found in a previous study as a result of selection for resistance to histidine of the Δ*dac* strain (11). It is therefore likely, that the *azlB* gene is inactivated in all suppressor mutants. The inactivation of *azlB* causes overexpression of the genes of the *azl* operon that are expressed downstream of *azlB* (10, 11). This involves the bipartite amino acid exporter AzlCD. To test the role of AzlB and AzlCD in the resistance to and export of L-Asn, we constructed strains carrying deletions of *azlB* and *azlBCD* and performed drop dilution assays with the resulting Δ*ansAB* Δ*ansZ* Δ*azlB* and Δ*ansAB* Δ*ansZ* Δ*azlBCD* triple mutants (GP4236 and GP4237, respectively) (Fig. 3). We observed improved growth when *azlB* was mutated or deleted, demonstrating that the inactivation of the *azlB* gene is responsible for the resistance to L-Asn. The additional deletion of the *azlCD* genes encoding the amino acid exporter resulted in the loss of growth. This finding suggests that AzlCD exports L-Asn in addition to 4-azaleucine and histidine (10, 11) (see Fig. 7).

### AimA is the major transporter for L-Asn on complex medium

The *B. subtilis* strain SP1 is a prototrophic derivative of the laboratory strain 168 that has been used throughout this study. Due to its prototrophy this strain is of interest for biotechnological applications (40). SP1 lacking the *ansAB* operon (strain BP269) is also sensitive to L-Asn, even on complex medium. Again, we were able to isolate suppressor mutants that could tolerate the addition of 19 mM L-Asn to LB medium. Genome sequencing of two suppressor mutants (see Table 1) revealed two distinct mutations that affect the amino acid importer AimA. In the mutant BP391, there was an insertion of three bases in the *aimA* gene resulting in the insertion of a Thr residue after Ser-64 in AimA. In the second mutant, BP392, there was a single nucleotide insertion in *aimA*, resulting in the expression of a truncated protein. In addition, BP391 had a second mutation that resulted in the substitution of Ala-96 by a Glu residue in the YetL protein, which acts as a transcriptional repressor to control the expression of the *yetM* (encodes a FAD-dependent monooxygenase) and *yetL* genes (41). Since these functions seem to be unrelated and the inactivation of AimA was already shown to be sufficient to cause resistance to L-Asn, the *yetL* mutation was not further investigated. In conclusion, the suppressor screen using the SP1-derived strain BP269 confirms the important role of AimA in the uptake of L-Asn.

As we have identified AimA as an importer for L-asparagine, we also deleted the *aimA* gene in the Δ*ansAB* Δ*ansZ* mutant in order to test whether the strain would still be sensitive to L-Asn stress on minimal medium. Interestingly, the deletion of the *aimA* gene only conferred limited resistance to L-asparagine (see Fig. 3, GP4239) if the bacteria grew on minimal medium. Again, this finding indicates that there are multiple transporters for L-Asn in *B. subtilis*.

### Further adaptation to L-Asn identifies BcaP as additional L-Asn importer

In order to increase the selective pressure, we employed a disc diffusion assay of a strain that lacks the main known options to adapt to the presence of L-Asn (the Δ*ansAB* Δ*ansZ* Δ*azlBCD* Δ*aimA* quadruple mutant GP4245) with high concentrations of L-Asn (500 mM) (see Fig. 4). We observed suppressor mutant formation after 48h of incubation at 42°C. We isolated a total of eight suppressors and sequenced the whole genomes of two of them. The results are summarized in Table 1. Both mutants carried mutations affecting the amino acid transporter BcaP. In strain GP4251, BcaP had a substitution of Ala-68 in Glu, and in GP4252, there was a base substitution that resulted in the direct formation of a stop codon and thus to a truncated BcaP protein. In addition, strain GP4251 had a second mutation in the *ydeC* gene that resulted in a substitution of His-43 to Gln in the AraC-type transcription factor YdeC of unknown function. We then sequenced the *bcaP* and *ydeC* alleles of the remaining six mutants. All had mutations in *bcaP*, while none of them had the mutation in *ydeC*. Therefore, we focussed the further analysis on *bcaP*.

**FIG 4.**
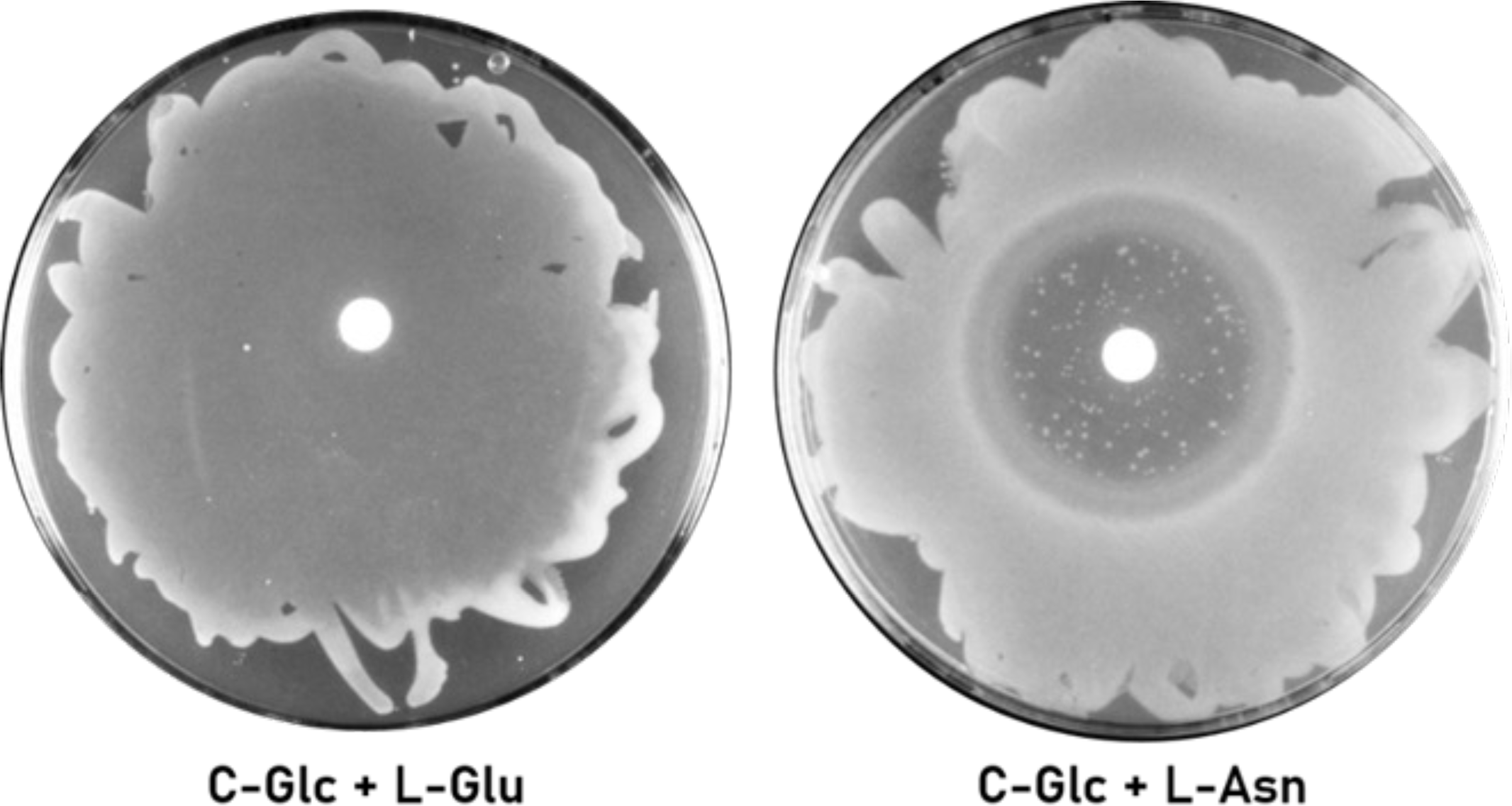
Disc diffusion assay of the Δ*ansA* Δ*ansZ* Δ*azlBCD* Δ*aimA* (GP4245) strain. 15 µl 150 mM L-glutamate solution was used as positive control and dropped on the filter disc. 15 µl 500 mM L-asparagine solution was used to apply the selective pressure. Suppressor mutants appeared after 48h of incubation at 42°C.

All eight suppressors carried mutations in *bcaP*, the gene coding for the branched chain amino acid permease BcaP, which is involved in the uptake of isoleucine, valine, threonine and serine (3, 5, 9). The mutations appear to make BcaP nonfunctional, which suggests a role of BcaP in the uptake of L-asparagine. When we compared our suppressors as well as the effect of a deletion of the *bcaP* gene (in strain GP4267) with the original strain (Δ*ansAB* Δ*ansZ* Δ*azlBCD* Δ*aimA*) in a drop dilution assay, we found that both the suppressor mutants as well as the *bcaP* deletion mutant are fully resistant to L-Asn stress (see Fig. 5). Thus, the lack of both AimA and BcaP confers full resistance to L-Asn, indicating that these two transporters are responsible for the uptake of this amino acid.

**FIG 5.**
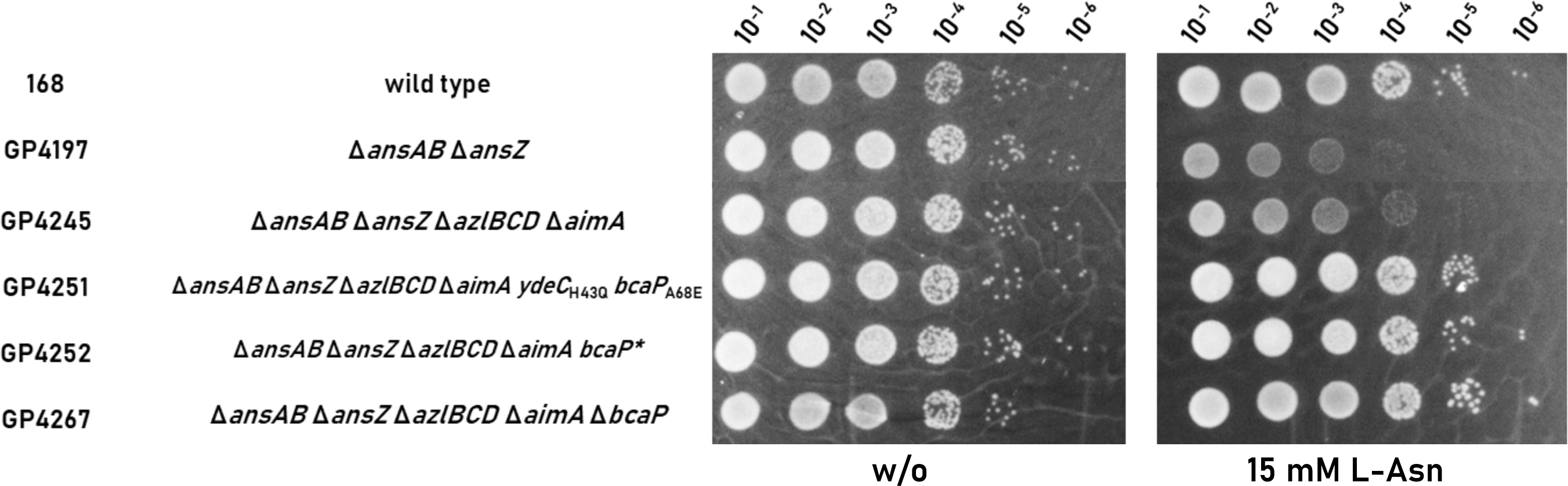
BcaP is also an Importer for L-Asn. Sensitivity of both suppressors (GP4251, GP4252) isolated via disc diffusion assay, as well as the Δ*ansAB* Δ*ansZ* Δ*azlBCD* Δ*aimA* Δ*bcaP* mutant (GP4267) to L-Asn is shown. The cells were grown in C-Glc minimal medium to an OD_600_ of 1.0 and were then diluted 10- fold to create dilutions ranging from 10^-1^ to 10^-6^. The dilution series was dropped onto C-Glc plates with and without L-asparagine (0 and 15mM) respectively. The plates were incubated at 42°C for 48 h.

The AimA amino acid importer and the AzlCD amino acid exporter have been shown to be involved in the uptake and export, respectively, of multiple amino acids. In order to characterize the suppressor mutants further, we also carried out an experiment with L-serine. L-Ser is toxic to *B. subtilis* in minimal medium (3, 15) To test, whether AimA, BcaP, and AzlCD are also involved in the homeostasis of L-Ser, we examined the growth of the wild type strain 168, the Δ*azlB* mutant GP3600, the Δ*ansAB* Δ*ansZ* Δ*azlBCD* Δ*aimA* mutant GP4245 and of the two suppressor mutants GP4251 and GP4252 at different concentrations of L-Ser (see Fig. 6). As observed before (3), the wild type strain was unable to grow if L-Ser was present. The same result was obtained for the *azlB* mutant GP3600 indicating that the AzlCD amino acid exporter does not play a significant role in serine export. At L-Ser of 5 mM, the loss of the amino acid importer AimA in GP4245 provided a very faint protection against growth inhibition by L-Ser. However, both suppressor mutants that are otherwise isogenic to the progenitor GP4245, exhibited a strong resistance to the presence of serine. This oberservation is in good agreement with previous reports that BcaP and AimA make the major contributions to L-Ser uptake. Interestingly, the two suppressor mutants differ in their resistance to L-Ser at higher concentrations. The *aimA bcaP* double mutant GP4252 barely grew at 15 mM serine or above. In contrast, GP4251 which has an additional mutation affecting the AraC family regulator YdeC grew well at 15 mM of L-Ser indicating that the mutation in *ydeC* contributes to amino acid resistance (see Discussion).

**FIG 6.**
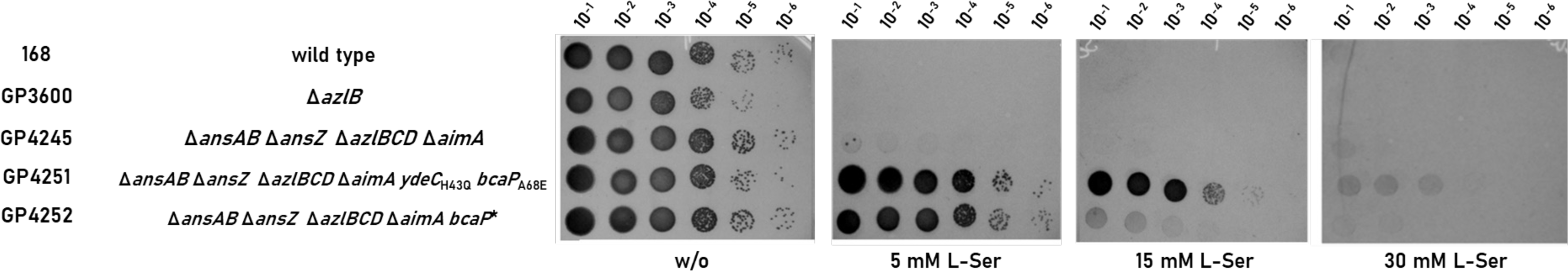
The isolated suppressors are also resistant against L-serine. Sensitivity of wild type *B. subtilis* (168), the Δ*azlB* mutant (GP3600), the Δ*ansAB* Δ*ansZ* Δ*azlBCD* Δ*aimA* mutant (GP4245) and the two suppressors isolated from L-Asn stress (GP4251 and GP4252) to L-serine is shown. The cells were grown in C-Glc minimal medium to an OD_600_ of 1.0 and were then diluted 10-fold to create dilutions ranging from 10^-1^ to 10^-6^. The dilution series was dropped onto C-Glc plates with and without L-serine (0, 5, 15 and 30 mM respectively) The plates were incubated at 42°C for 48 h.

**FIG 7.**
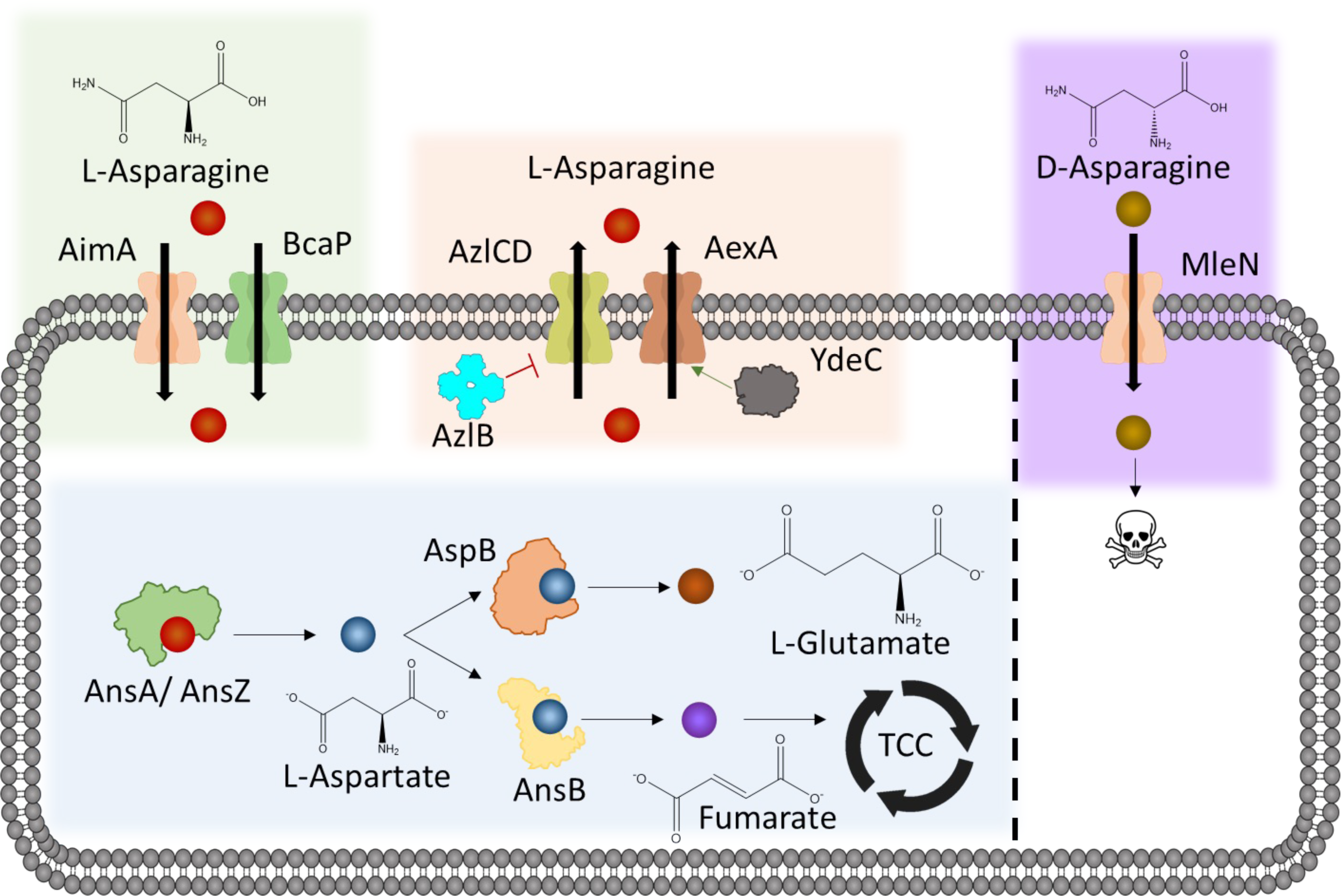
Asparagine metabolism in *B. subtilis*. L-asparagine enters the cell via the importers AimA and BcaP. Within the cell it is degraded to L-aspartate and then subsequently feeds into the tricarboxylic acid cycle (TCC) and production of L-glutamate via a newly discovered AspB-dependent bypass (59). L-asparagine is exported from the cell due to mutations in the regulators AzlB and YdeC. Export happens by the broad range amino acid exporters AexA (YdeD) and AzlCD. D-asparagine enters the cell via MleN and is toxic to *B. subtilis*. No form of degradation or export was detected.

## DISCUSSION

This work aimed to utilize the toxic effects of D-asparagine to identify uptake mechanisms for both D- and L-asparagine, following the hypothesis that both enantiomers enter the cell via the same transporter, as it is the case for alanine in *B. subtilis* (12). The mutations in MleN revealed it to be a D-Asn importer. Even in the presence of MleN, L-Asn was well tolerated if the two transporters AimA and BcaP were missing indicating that MleN is not involved in the uptake of L-Asn. This demonstrates that, although structurally similar, D- and L-asparagine are taken up by different uptake systems. Our suppressor screen didn’t give any indication of the possiblity of D-Asn export in *B. subtilis*. This is interesting since we were easily able to find mutations in the *azlB* repressor gene that result in consitutive expression of the amino acid exporter AzlCD in the presence of L-Asn. This observation again supports the idea, that even though amino acid transporters are often quite promiscous (2), it can not be taken for granted that transporters that import or export any given amino acid do so for the L- and D-forms of the amino acid. It should be noted that the high concentration of D-Asn used in the suppressor screen does not reflect physiological conditions. Therefore, it cannot be ruled out that *B. subtilis* encodes additional high-affinity transporters for D-Asn that might be active at lower concentrations.

A critical point for the identification of bacterial amino acid transporters is the use of strains and/or conditions that make the target amino acid toxic for the cells. In this study, we used strains lacking the second messenger c-di-AMP (Δ*dac*) and strains that are unable to degrade asparagine (Δ*ansAB* mutant) resulting in the intracellular accumulation of L-Asn. Interestingly, the different strains seem to have experienced different selective pressures resulting in the identification of highly specific suppressor mutations according to the selection scheme. For the Δ*dac* mutant, the well- established sensitivity to potassium was the major bottleneck, resulting in the isolation of *ktrD* mutants in all four cases (see below). For the strain lacking both asparaginases, the intracellular accumulation of L-Asn was the main issue which could be efficiently solved by activating the AzlCD amino acid exporter in all four independently isolated strains. Interestingly, the prototrophic strain SP1 lacking the asparaginase AnsA inactivated the AimA amino acid importer rather than activating AzlCD. This might result from the use of complex medium in which several amino acids share transporters. Moreover, the expression of the second transporter BcaP is repressed during growth on complex medium by the transcription factor CodY (42). Thus, under these conditions, AimA might be the only relevant transporter for L-Asn, resulting in its preferred inactivation to cope with L-Asn stress. Finally, when degradation or export of L-Asn were blocked, and one of the uptake systems (AimA) was also missing, mutations in a second transporter BcaP were selected in all eight cases. Thus, our results highlight the specificity of the outcomes that result from even subtle differences in the selective scenario.

In the case of the Δ*dac* mutant GP2222, all suppressor mutants had the KtrD potassium channel inactivated. This protein imports potassium with low affinity, but it has a very high affinity for the ion in the presence of glutamate (4). It can therefore be concluded that the well-established potassium sensitivity of the Δ*dac* mutant (4, 18, 43) is the major factor that causes growth inhibition in the presence of L-Asn, resulting in mutations that inactivate the strongly expressed KtrD potassium channel. Similarly, a mutation in *ktrD* was also selected if the Δ*dac* mutant grew in the presence of high concentrations of histidine (11). While glutamate is one one the final degradation products of histidine, no glutamate is formed during the utilization of L-Asn. This suggests that either L-Asn or its degradation product L-Asp can also cause an increased affinity of KtrD for potassium, as shown for glutamate (4).

Our previous work has shown that the AimA protein is a major non-specific player in amino acid uptake (3, 4). Based on our results with both the Δ*dac* Δ*aimA* mutant GP3054 (see Fig. 2) and the SP1 Δ*ansAB* mutant BP269 (see Table 1), AimA also plays the major role in the uptake of L-Asn. Thus, AimA is the major importer for glutamate, serine and also asparagine (3, 4). It is tempting to speculate that AimA might also be involved in the transport of other amino acids. Since AimA was discovered only recently, its presence may have masked previous genetic attempts to identify amino acid transporters in *B. subtilis*. AimA is a member of the large amino acid-polyamine-organocation superfamily (44). These proteins typically transport amino acids, but they may also be involved in the uptake of methyl-thioribose (MtrA) or potassium ions (KimA) (18, 45). Interestingly, AimA seems to be limited to *B. subtilis*, as close orthologs of the protein are missing in most species, even in other *Bacillus* species. In *B. subtilis*, AimA has a paralog, YveA. This protein is expressed during sporulation in the forespore (46) and may thus be important for amino acid uptake once the spores start to germinate. However, based on the expression profile, YveA is unlikely to contribute to amino acid transport in growing cell.

Based on the suppressor mutants, the selective pressure caused by L-Asn was different in strains that are unable to degrade this amino acid from those that lack c-di-AMP. In the case of the 168-derived *ansAB ansZ* mutant all mutations affected the AzlB transcription repressor. Such mutations were previously shown to cause expression of the amino acid exporter complex AzlCD and to allow export of toxic histidine (11). Moreover, an amino acid export activity of AzlCD was also suggested for 4-azaleucine (10). Indeed, the loss of the exporter subunits in addition to the repressor results in an increased sensitivity towards L-Asn as compared to the *azlB* repressor mutant (see Fig. 3). This suggests that the AzlCD amino acid exporter is responsible for the resistance to L-Asn by exporting this amino acid. However, we cannot exclude the possibility that the accumulation of L-Asn in the *ansAB ansZ* mutant results in the formation of toxic products that are the actual substrate for export by AzlCD. Taking into account that AzlCD is a member of a family of amino acid exporters (47), and all the observations of this study it seems most likely that L-Asn is indeed the substrate for AzlCD.It is interesting to note that L-Asn is already the third amino acid that is likely to be exported by the AzlCD exporter. Moreover, expression of AzlCD also provides resistance to otherwise toxic diaminopropionic acid (Warneke et al., unpublished results), thus suggesting that this complex is a broad-spectrum amino acid exporter. So far, no conditions have been identified that would allow a substantial expression of the AzlCD amino acid exporter (35), and the acquisition of resistance to growth-inhibiting amino acids always depends on the inactivation of the *azlB* repressor gene. It is tempting to speculate that the untimely expression of the amino acid exporter might cause a loss of amino acids from the cell. Since the bacteria invest a lot of energy to either synthesize or import amino acids, their loss would be disadvantage for the bacteria. Indeed, the fitness of the *azlB* mutant is reduced in minimal media with ammonium as the single source of nitrogen, when the cells depend on de novo biosynthesis of amino acids (48). Thus, it might be a good strategy to encode the exporter in the genome but to keep the gene silent unless expression is required because a toxic amino acid causes the corresponding selective pressure.

As stated above, the identification of conditions that render an amino acid toxic is crucial to find the corresponding transport systems. Another step in this strategy is to use mutants that are unable to activate *e.g.* amino acid exporters or that are already defective in main transporters. Indeed, this approach allowed us to identify BcaP as a second importer for L-Asn. As for AimA, several substrates have been identified for BcaP. BcaP is the major transporter for the branched-chain amino acids isoleucine and valine as well as for threonine (3, 5, 9). Moreover, in addition to AimA, BsaP plays a minor role in serine uptake (3). The observation that BcaP is also involved in the acquisition of L-Asn suggests that it is also a more non-specific amino acid importer.

In one of the suppressor mutants that had acquired the mutations in *bcaP*, we found a second mutation in the *ydeC* gene encoding an unknown transcription regulator of the AraC family. It is tempting to speculate that YdeC controls the expression of an amino acid transporter. The observation that the presence of the *ydeC* mutation causes an increased resistance towards serine suggests again a general role for the corresponding transporter. Indeed, this transporter, AexA (previously YdeD), can export β-alanine (Warneke et al., unpublished results) and probably also L-Asn and L-Ser.

The work described here has identified systems for the uptake and export of D- and L-Asn in *B. subtilis* (see Fig. 7). As for many other proteinogenic amino acids, L-Asn can be imported and exported by multiple broad-range transport proteins.

## MATERIALS AND METHODS

### Strains, media and growth conditions

*E. coli* DH5α was used for cloning. All *B. subtilis* strains used in this study are derivatives of the laboratory strains 168 or SP1. *B. subtilis* and *E. coli* were grown in Luria-Bertani (LB) or in sporulation (SP) medium (49, 50). For growth assays and the *in vivo* interaction experiments, *B. subtilis* was cultivated in LB, SM, MSSM or C-Glc minimal medium (18). SM is a minimal medium that uses KH_2_PO_4_ as buffer. MSSM is a modified SM medium in which KH_2_PO_4_ was replaced by NaH_2_PO_4_ and KCl was added as indicated. C-Glc is a chemically defined medium that contains glucose (1 g/l) as carbon source (51). The media were supplemented with ampicillin (100 µg/ml), kanamycin (10 µg/ml), chloramphenicol (5 µg/ml), spectinomycin (150 µg/ml), tetracycline (12.5 µg/ml) or erythromycin plus lincomycin (2 and 25 µg/ml, respectively) if required.

### DNA manipulation and transformation

Transformation of *E. coli* and plasmid DNA extraction were performed using standard procedures (49). All commercially available plasmids, restriction enzymes, T4 DNA ligase and DNA polymerases were used as recommended by the manufacturers. *B. subtilis* was transformed with plasmids, genomic DNA or PCR products according to the two-step protocol (50). DNA fragments were purified using the QIAquick PCR Purification Kit (Qiagen, Hilden, Germany). DNA sequences were determined by the dideoxy chain termination method (49).

### Construction of mutant strains by allelic replacement

Deletion of the *ansAB, aimA* and *bcaP* genes was achieved by transformation of *B. subtilis* 168 with a PCR product constructed using oligonucleotides to amplify DNA fragments flanking the target genes and an appropriate intervening resistance cassette as described previously (52). The integrity of the regions flanking the integrated resistance cassette was verified by sequencing PCR products of about 1,100 bp amplified from chromosomal DNA of the resulting mutant strains. In the case of *ansAB*, the cassette carrying the resistance gene lacked a transcription terminator to ensure the expression of the downstream genes.

### Phenotypic analysis

In *B. subtilis*, amylase activity was detected after growth on plates containing nutrient broth (7.5 g/l), 17 g Bacto agar/l (Difco) and 5 g hydrolyzed starch/l (Connaught). Starch degradation was detected by sublimating iodine onto the plates.

Quantitative studies of *lacZ* expression in *B. subtilis* were performed as follows: cells were grown in C-Glc medium. Cells were harvested at OD_600_ of 0.5 to 0.8. β-Galactosidase specific activities were determined with cell extracts obtained by lysozyme treatment as described previously (50). One unit of β-galactosidase is defined as the amount of enzyme which produces 1 nmol of o-nitrophenol per min at 28° C. All β-galactosidase assays were performed in triplicate.

To assay the growth of *B. subtilis* mutants at different asparagine concentrations, multiple drop dilution assays was performed. Briefly, precultures in either MSSM, C-Glc or SM minimal medium at the indicated L- and D-asparagine concentration were washed three times and resuspended to an OD_600_ of 1.0 in a 1 x MSSM buffer, C-salts or SM-buffer solution. A dilution series was then pipetted onto the respective minimal medium plates containing the desired asparagine concentration. All drop dilution assays were performed in triplicate.

### Disc Diffusion Assay

For the disc diffusion assay a preculture of GP4245 in C-Glc medium was prepared. The cells were harvested during exponential growth phase, washed three times and then resuspended to an OD_600_ of 1.0 in a 1 x C-salts solution. Then, 150µl of the cells were spread onto C-Glc agar plates and left for 2 minutes to dry. A sterile filter disc was placed in the middle of the agar plates and 15µl of highly concentrated (500mM) L-asparagine solution was then pipetted onto the filter disc. A plate with 150mM L-glutamate solution was used as positive control. The plates were incubated at 42°C for 48 h and then photographed and the suppressors isolated.

### Genome sequencing

To identify the mutations in the suppressor mutant strains BP391, BP392, GP4158, GP4159, GP4230, GP4231, GP4232, GP4233, GP4251, GP4252, GP4269, GP4270 (see Tables 1, 3), the genomic DNA was subjected to whole-genome sequencing. Concentration and purity of the isolated DNA was first checked with a Nanodrop ND-1000 (PeqLab Erlangen, Germany), and the precise concentration was determined using the Qubit® dsDNA HS Assay Kit as recommended by the manufacturer (Life Technologies GmbH, Darmstadt, Germany). Illumina shotgun libraries were prepared using the Nextera XT DNA Sample Preparation Kit and subsequently sequenced on a MiSeq system with the reagent kit v3 with 600 cycles (Illumina, San Diego, CA, USA) as recommended by the manufacturer. The reads were mapped on the reference genome of *B. subtilis* 168 (GenBank accession number: NC_000964) (53). Mapping of the reads was performed using the Geneious software package (Biomatters Ltd., New Zealand) (54). Frequently occurring hitchhiker mutations (55) and silent mutations were omitted from the screen. The resulting genome sequences were compared to that of our in-house wild type strain. Single nucleotide polymorphisms were considered as significant when the total coverage depth exceeded 25 reads with a variant frequency of ≥90%. All identified mutations were verified by PCR amplification and Sanger sequencing.

**TABLE 3.**
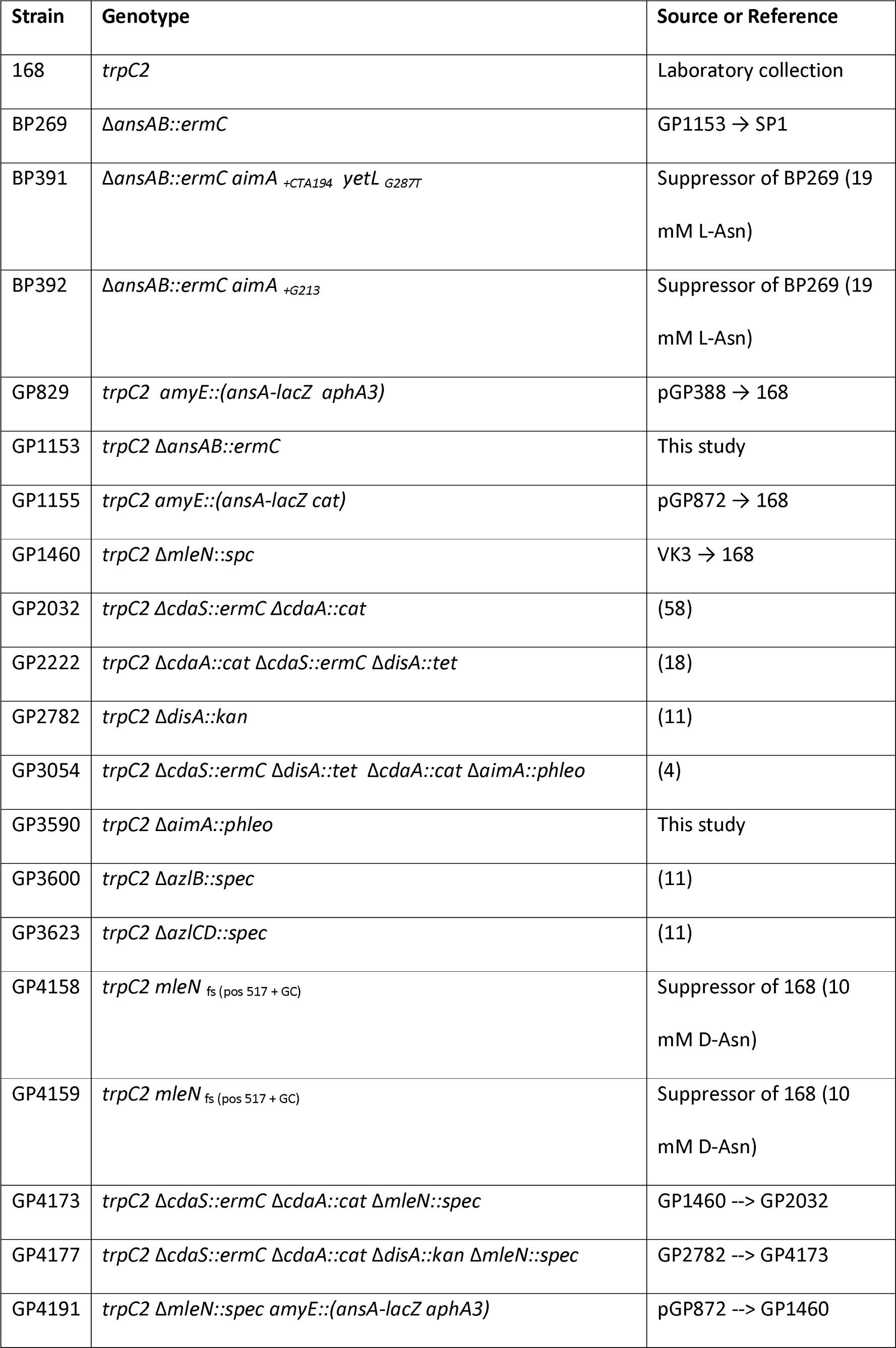
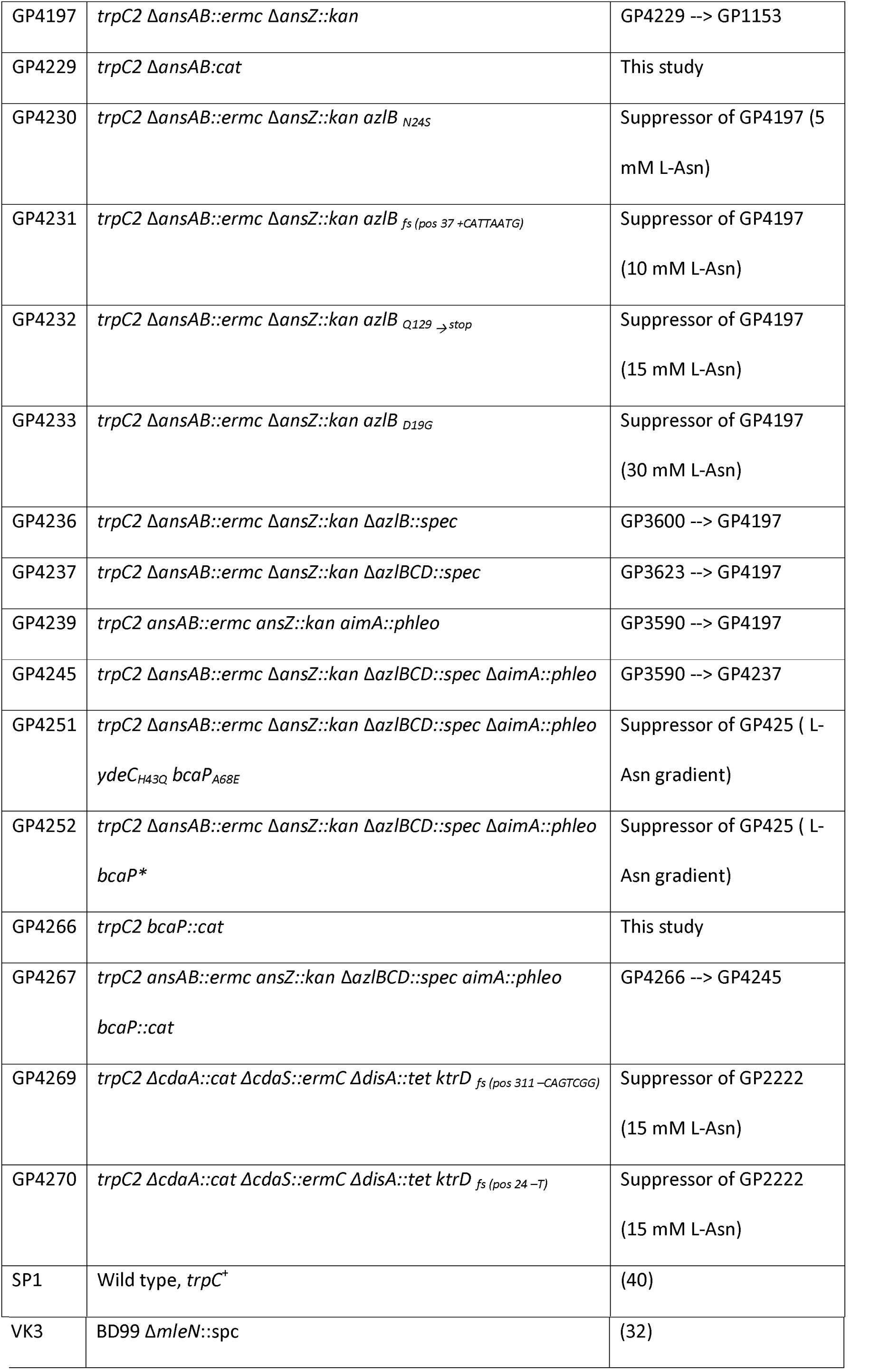
*B. subtilis* strains used in this study.

### Plasmid construction

To construct translational fusions of the potential *mleN* and *ansA* promoter regions to the promoterless *lacZ* gene, we used the plasmids pAC7 (56) and pAC5 (57), respectively. Briefly, the promoter regions were amplified using oligonucleotides that attached EcoRI and BamHI restriction to the ends of the products, and the fragments were cloned between the EcoRI and BamHI sites of pAC5 or pAC7. The resulting plasmids were pGP388 for *mleN* and pGP872 for *ansA*, respectively.

## ACKNOWLEDGEMENTS

We are grateful to Jens Landmann, Paul Molis, and Aneta Zelazo for the help with some experiments. This work was supported by grants from the Deutsche Forschungsgemeinschaft via Priority Programme SPP1879 (to J.S. and F.M.C.).

